# Selecting for *Chlamydomonas reinhardtii* fitness in a liquid algal growth system compatible with the International Space Station Veggie plant growth chamber

**DOI:** 10.1101/2020.01.13.904342

**Authors:** Junya Zhang, Bárbara S.F. Müller, Kevin N. Tyre, Fang Bai, Ying Hu, Marcio F.R. Resende, Bala Rathinasabapathi, A. Mark Settles

**Author notes:** **Correspondence:** A. Mark Settles.

## Abstract

A biological life support system for spaceflight would capture carbon dioxide waste produced by living and working in space to generate useful organic compounds. Photosynthesis is the primary mechanism to fix carbon into organic molecules. Microalgae are highly efficient at converting light, water, and carbon dioxide into biomass, particularly under limiting, artificial light conditions that are a necessity in space photosynthetic production. Although there is great promise in developing algae for chemical or food production in space, most spaceflight algae growth studies have been conducted on solid agar-media to avoid handling liquids in microgravity. Here we report that breathable plastic tissue culture bags can support robust growth of *Chlamydomonas reinhardtii* in the Veggie plant growth chamber, which is used on the International Space Station to grow terrestrial plants. Live cultures can be stored for at least one month in the bags at room temperature. The gene set required for growth in these photobioreactors was tested through a short-wave ultraviolet light (UVC) mutagenesis and selection experiment with wild-type (CC-5082) and *cw15* mutant (CC-1883) strains. Genome sequencing identified UVC-induced mutations, which were enriched for transversions and nonsynonymous mutations relative to natural variants among laboratory strains. Genes with mutations indicating positive selection were enriched for information processing genes related to DNA repair, RNA processing, translation, cytoskeletal motors, kinases, and ABC transporters. These data suggest modification of signal transduction and metabolite transport may be needed to improve growth rates in this spaceflight production system.

## 1 Introduction

Microalgae grow by converting light, water, and CO_2_ into biomass. Algae have long been proposed for space life support systems to recycle CO_2_ and provide food either directly or indirectly to astronauts (Brechignac and Schiller, 1992; Ai et al. 2008; Niederwieser et al., 2018; Matula and Nabity, 2019). Many species of microalgae are photosynthetically efficient under the limiting light and low volume conditions necessary in space production (Kliphuis et al., 2012). As single cell organisms, microalgae are easy to cultivate with minimal requirements, and have great potential to yield value-added products.

Both eukaryotic and prokaryotic species, such as *Chlorella vulgaris* and *Arthrospira platenis*, respectively are generally regarded as safe (GRAS) for human consumption (Caporgno and Mathys, 2018). Algae have nutritional benefits with high levels of antioxidants and protein with essential amino acids (Buono et al., 2014). Algae are also rich in ω-3 fatty acids, such as eicosapentaenoic acid and docosahexaenoic acid (Salem and Eggersdorfer, 2015). The red algal carotenoid, astaxanthin, has multiple uses including the ability to protect against retinal damage in animals (Wang et al., 2000; Katagiri et al., 2012; Liu et al., 2016; Otsuka et al., 2016; Shah et al., 2016). Algal oils may be used in future astronaut diets to help mitigate harmful effects of microgravity and cosmic radiation during spaceflight.

A few past studies exposed or grew algal species in space conditions (Mergenhagen and Mergenhagen, 1989; Wang et al., 2006; Giardi et al., 2013; Niederwieser et al., 2018). A Space Shuttle mission in 1985 used *Chlamydomonas reinhardtii* to examine phototaxic responses in microgravity (Mergenhagen and Mergenhagen, 1989). Microgravity allowed cells to remain close to light sources in spaceflight suggesting that photosynthetic productivity could be higher in space. By contrast, Wang et al. (2006) grew cyanobacteria for 15 days in a satellite and found reduced growth relative to a ground control. However, the ground control temperature and light cycles were not adjusted to the space conditions. The satellite had two temperature drops and two missed day photoperiods preventing strong conclusions from being drawn about growth rates.

Giardi et al. (2013) compared Chlamydomonas photosynthetic responses for wild-type and photosystem II (PSII) D1 mutants in stationary cultures. Spaceflight had a greater negative effect on PSII fluorescence in wild-type cells and negatively impacted cell growth upon return to Earth. Soviet and Russian spaceflight experiments indicate that *Chlorella vulgaris* has similar growth kinetics in space and on Earth, but the cells experience spaceflight as a stress (reviewed in Niederwieser et al., 2018).

The impact of the spaceflight environment for larger scale production of algae is currently unknown. To test larger scale microalgae production, the European Space Agency (ESA) has developed a photobioreactor that pumps liquid media through a meandering path, pipe-reactor (Bretschneider et al., 2016). Gas exchange is mediated by fluorinated ethylene propylene (FEP) membranes that allow CO_2_ and O_2_ diffusion, while algae are continuously mixed with a peristaltic pump (Helisch et al., 2019). This photobioreactor is currently being tested on the International Space Station (ISS) for long-term growth of *Chlorella vulgaris.* A significant challenge of maintaining photosynthetic productivity is regular removal of stationary cells and addition of new media to maintain photoautotrophic growth without forming excessive biofilms within the raceway. In addition, this photobioreactor experiment is not focusing on adapting algae species to spaceflight conditions (Helisch et al., 2019).

In yeast, competitive growth experiments in liquid culture were used to identify genes needed for survival in spaceflight conditions (Nislow et al., 2015). We are completing spaceflight experiments to determine genes required for growth of Chlamydomonas via competitive growth of mutagenized cells. Here we report results from our development of methods and experiment verification test (EVT). We describe a simple batch culture protocol using commercial FEP tissue culture bags that is adapted to the Veggie plant growth chambers on the ISS. Shortwave ultraviolet (UVC) light mutagenesis and full genome sequencing enabled the detection of new mutations in two microalgae strains. The spectrum of mutations identified suggest that UVC primarily induces DNA damage in Chlamydomonas via errors in translesion synthesis and double strand break repair. In addition, a variety of cellular information processing functions may need to be modified to improve growth rates in this batch culture system.

## 2 Materials and Methods

### 2.1 Strains and culturing conditions

Strains CC-5082 (WT) and CC-1883 (*cw15*) were obtained from the Chlamydomonas Resource Center. CC-5082 is a sequence-verified clone of CC-1690, which is a wild-type strain of mating type *mt*^*+*^ (Gallaher et al., 2015). CC-1883 is an *mt*^-^ *cw15* cell wall mutant, which allows transformation of exogenous DNA by vortexing with cells and glass beads (Kindle et al., 1989). Strains were maintained at room temperature (20-25°C) on agar plates with Tris acetate-phosphate (TAP) medium (Gorman and Levine, 1965), under 50 to 100 μmol/m^2^/s continuous photosynthetically active radiation (PAR) from daylight fluorescent bulbs (6500 K).

To initiate liquid cultures, 1-2 mm colonies were scraped from TAP agar plate and inoculated into 50 mL TAP media. Traditional liquid cultures were grown with continuous light in 250 mL Erlenmeyer flasks with 100 rpm gyratory shaking. For spaceflight analogs, liquid cultures were grown in 120 mL PermaLife cell culture bags (OriGen Biomedical, Austin, TX, USA). Liquid cultures were grown in daylight fluorescent lighting, LED lighting with a ratio of red: green: blue of 4:1:1, or in the Veggie Vegetable Production System at the Kennedy Space Center (Merritt Island, FL, USA) with a ratio of red: green: blue of 8:1:1. All lighting was 80-100 μmol/m^2^/s.

The UVC mutagenesis dose that caused ~10% cell survival was determined by transferring 7 mL of early-log phase culture at OD_600_ = 0.45 to 15 cm sterile petri plates in a sterile laminar flow hood. The petri plates were opened in a GS Gene Linker UV Chamber (Bio-Rad, Hercules, CA, USA) and exposed to increasing doses of UVC light from germicidal bulbs. Petri plates were closed, wrapped in aluminum foil, and agitated overnight in the dark at 50 rpm. Mutagenized cultures were plated on TAP agar plates with non-treated samples diluted 1:5 prior to plating. Colonies were grown under continuous light, counted, and normalized relative to non-mutagenized cultures.

### 2.2 EVT mutagenesis

Colonies from TAP agar plates were scraped and suspended in 600 μL liquid TAP media and adjusted to an optical density of 1 at 600 nm using a visible light spectrophotometer (SmartSpec 3000, Bio-Rad, Hercules, CA, USA). The Chlamydomonas suspension was used to inoculate TAP liquid media at a 1:250 dilution, i.e. 0.2 mL of OD_600_=1 suspension was added to 50 mL TAP media. WT was inoculated on day 1 of the experiment and *cw15* was inoculated on day 2. The cultures were grown in flasks until reaching an OD_600_ of 0.4-0.5 on day 4. Non-mutagenized cells from the culture were sampled for whole genome sequencing by centrifuging 2 mL of the culture and freezing the cell pellet at −80°C until DNA was extracted. For each mutagenesis, 7 mL of early-log phase culture was exposed to 8 mJ of UV light. The mutagenized cells were then used to inoculate 100 mL of TAP media in a PermaLife tissue culture bag in a sterile laminar flow hood. Inoculated tissue culture bags were held at room temperature in the dark for 7 d and then transferred to the Veggie growth chamber.

### 2.3 EVT culture conditions and selection

The Veggie chamber was set with the reservoir at maximum distance from the lighting. The red light was set to ‘medium’; the blue light was set to ‘low’; and the green light was set to ‘on’. The bellows were closed during growth cycles, and the fan was set to ‘low’. The initial mutagenized cultures were grown for 7 d in Veggie. The cultures were then passaged by transferring 1 mL of culture to a bag containing fresh TAP media using a sterile syringe. The second culture was grown for 6 d and then passaged to a third tissue culture bag for 6 d of growth. During each passage, 2 mL of culture was sampled, centrifuged, and the cell pellet was frozen at −80°C until DNA was extracted. The remaining cultures were stored in a soft stowage, Cargo Transport Bag (CTB) until 36 d after the initial inoculation. For each dark-stored culture, 2 mL was sampled for DNA extraction.

### 2.4 Biomass measurement

Dry biomass was determined by transferring all of the remaining culture volume into two 50 mL centrifuge tubes. The cells were centrifuged with the supernatant media remove and the cell pellets were lyophilized overnight. Dry weights were measured on an analytical balance. Biomass for each culture bag is the average of the two technical replicates.

### 2.5 Whole genome sequencing

DNA was extracted from the flash-frozen and dark-stored cell pellets as described with some minor modifications (Newman et al., 1990). Cell pellets were resuspended in 150 μL H_2_O on ice, and 300 μL of SDS-EB buffer (2% SDS, 400 mM NaCl, 40 mM EDTA, 100 mM Tris-HCl, pH 8.0) was added and vortexed. The cell suspension was then extracted with 350 μL phenol: chloroform: isoamyl alcohol 25:24:1 (v:v) for 2-5 min by intermittent mixing using a vortex. The organic and aqueous phases were separated with a 5 min centrifugation at maximum speed in a microcentrifuge. The aqueous phase was then extracted with 300 μL chloroform: isoamyl alcohol (24:1). The RNA in the aqueous phase was digested with RNase A at room temperature for 10 min. DNA was precipitated by adding 1 mL 100% ethanol, incubating on ice for 30 min, and centrifuging for 10 min. The pellet was washed with 200 μL 70% ethanol, dried in vacuum centrifuge and resuspended in 30 μL H_2_O. DNA sample integrity was evaluated with an Agilent TapeStation (Santa Clara, CA, USA), and concentration was determined with Qubit dsDNA HS Assay Kit (Life Technologies, Carlsbad, CA) according to the manufacturer’s instructions.

For each library, 3 μg of DNA was sheared to 350 bp average size using an S220 Focused-ultrasonicator (Covaris, Woburn, MA, USA). Barcoded, TruSeq PCR-Free Low Throughput libraries were prepared for each DNA sample following the manufacturer’s instructions (Illumina, San Diego, CA, USA). Each sample had a unique index from Illumina TruSeq DNA CD Indexes (96 indexes/samples). The libraries were split into two pools to sequence a total of 38 samples: 9 dark-stored and 9 flash-frozen samples for each strain, growth cycle, and biological replicate as well as the two non-mutagenized initial cultures. Library pools were color-balanced and sequenced with paired-end 150 bp reads on the Illumina HiSeq X platform at MedGenome (Foster City, CA, USA).

### 2.6 Read processing and genome alignment

The raw sequence reads were filtered using Trimmomatic v.0.38 to remove barcode and adaptor sequences in paired-end mode (Bolger et al., 2014). High quality reads were aligned to the *Chlamydomonas reinhardtii* reference genome (Merchant et al., 2007), v.5.6 from Phytozome, using BWA-mem (Li, 2013). Duplicate reads were removed using Picard MarkDuplicates, v.2.19.1 (http://broadinstitute.github.io/picard/) with default parameters. Reads near insertion-deletion polymorphisms (indels) were realigned using GATK v.3.8.1 RealignerTargetCreator and IndelRealigner (DePristo et al., 2011; McKenna et al., 2010).

### 2.7 Identifying sequence variants

Three software packages for detecting single nucleotide polymorphisms (SNPs) and small indels were compared. FreeBayes v.1.2.0 (Garrison and Marth, 2016) was implemented with the pooled continuous option. LoFreq v. 2.1.3.1 used default parameters (Wilm et al., 2012), while CRISP pooled, discrete parameters were: --poolsize 5 --perms 100000 --mmq 10 --minc 3 (Bansal, 2010; Bansal et al., 2011). FreeBayes called more variants for libraries with the lowest depth giving a negative correlation between the number of mapped reads and number of variants detected. LoFreq is restricted to calling variants for single samples and not detecting variants at a population level. Consequently, CRISP was determined to be the optimal approach for variant detection.

For each strain, CRISP genotyping results were filtered to remove variants with a call rate below 70% (i.e. missing data ≥30%) and monomorphic variants with a minor allele frequency (MAF) of zero. Monomorphic variants revealed non-mutated, natural variants relative to the reference genome. In addition, variants with low depth and low-quality mapped reads were removed based on CRISP assignments of “LowDepth, LowMQ10 and LowMQ20.” Finally, novel variants were scored by identifying exact matches to natural variants reported from whole genome sequencing of 39 Chlamydomonas laboratory strains (Gallaher et al., 2015).

### 2.8 Determining protein coding variants and selection

The effect of each variant on protein coding regions was predicted using SnpEff v.4.3t (Cingolani et al., 2012). Synonymous and nonsynonymous mutations from the primary SnpEff call were used to estimate the overall impact of UVC mutagenesis on protein coding genes. Synonymous (π_S_) and nonsynonymous (π_N_) nucleotide diversity was estimated for each gene using SNPGenie (Nelson et al., 2015). SNPGenie was run for all high-quality and novel variants independently, using both the forward strand and reverse complement strand. Positive selection was inferred when π_N_ > π_S_, and purifying selection was inferred when π_N_ < π_S_. Genes showing positive or purifying selection were tested for gene ontology (GO) term enrichment using AgriGO v2.0 using default parameters except for the “Hochberg (FDR)” multi-test adjustment method for the default false discovery rate of 0.05 (Tian et al., 2017).

## 3 Results

We tested commercial FEP tissue culture bags for the ability to support microalgae growth without agitation or active mixing of gases with liquids. Under LED lighting, the bags are able to support robust growth (Figure 1A-B). Time courses of growth show a 2 d delay in the bags with log phase between 4-6 d and stationary phase at 8 d for both WT and *cw15* strains (Figure 1C-D). These results indicate that FEP tissue culture bags provide sufficient gas exchange to support microalgae, but that Chlamydomonas laboratory strains are better adapted to grow in flasks.

**Figure 1.**
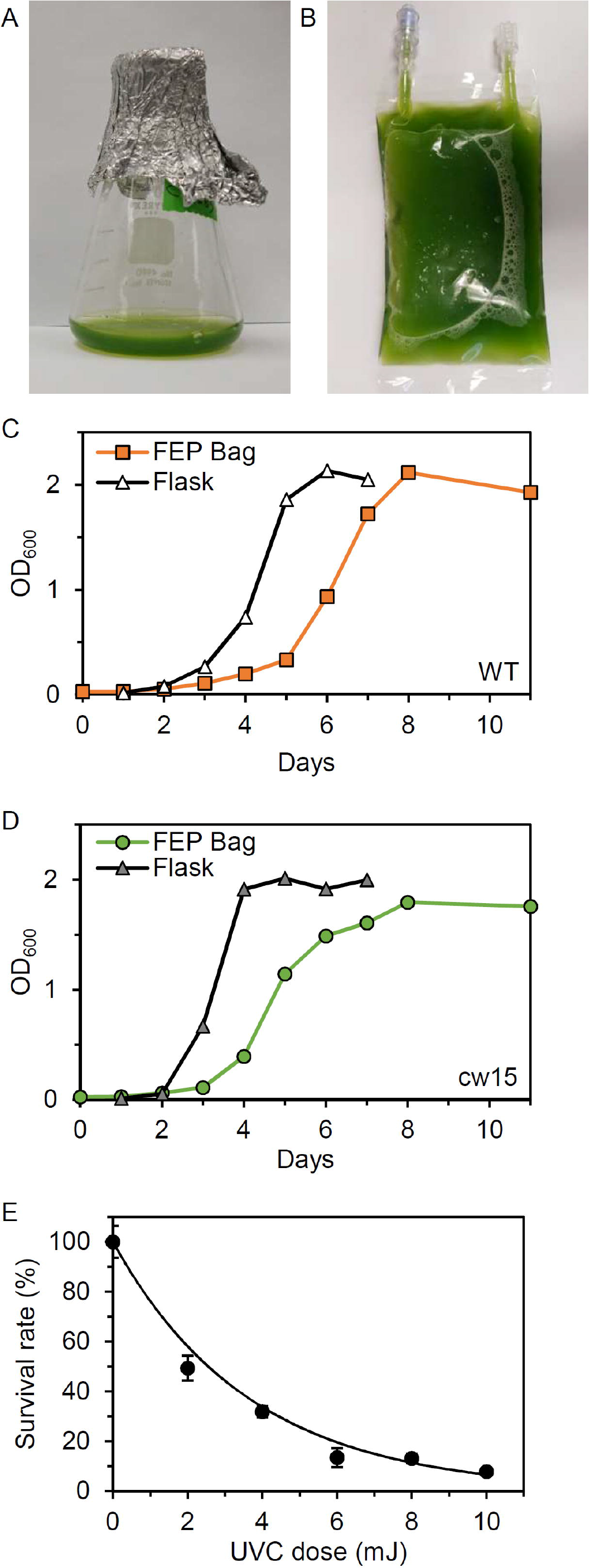
Microalgae growth in FEP plastic culture bags. (A) Flask culture of wild-type (WT), CC-5082. (B) FEP tissue culture bag of WT. (C) Growth curve of WT algae comparing growth in flasks with aeration and in FEP bags. (D) Growth curve of *cw15* (CC-1883) algae in flasks with aeration and in FEP bags. (E) Dose-response of WT algae to UVC light.

To select for mutations that would improve growth in tissue culture bags, we determined the dose-response for cell lethality in a UVC light chamber. Figure 1E shows that 6-10 mJ of UVC exposure is sufficient to kill ~90% of WT cells. Similar results were obtained for *cw15*, and we concluded that 8 mJ of UVC would give sufficient DNA damage to induce mutations in both strains without risking excessive cell death and culture failure during spaceflight.

An EVT was completed at the Kennedy Space Center (Figure 2). Three biological replicates of WT and *cw15* were mutagenized at the University of Florida, Gainesville, FL, USA. The mutagenized cells were transferred to tissue culture bags with TAP media and stored at room temperature in the dark for 7 d to emulate a late load and storage during a resupply mission to the ISS. The cultures were then transferred to a Veggie unit to provide light and stimulate photosynthesis. The culture bags were placed under bungee cords that hold plant pillows in the Veggie reservoir (Figure 2B). The bags were left without any agitation except during passages (Figure 2C). To complete a passage, culture bags were removed from the reservoir, agitated manually, and 1 mL of culture was transferred to fresh media for a new growth cycle (Figure 2D-F). An additional 2 mL of culture was sampled and cell pellets were frozen to preserve a DNA sample without dark storage.

**Figure 2.**
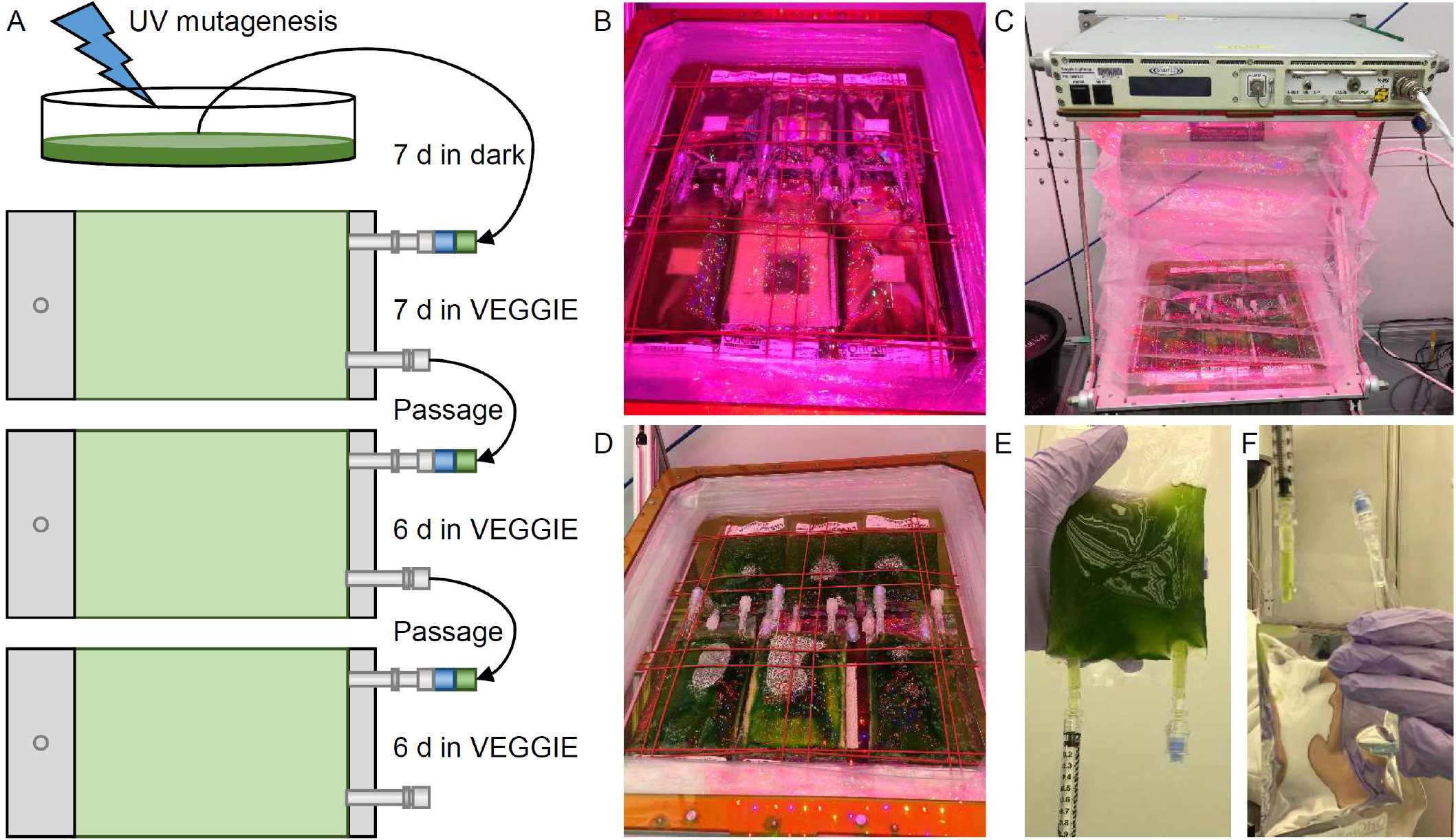
Algae selection experiment in Kennedy Space Center Veggie growth chamber. (A) Schematic of the experiment design and workflow. (B) Initial installation of mutagenized culture bags in the Veggie reservoir. (C) Veggie chamber with algae bags and closed bellows. (D) Algae cultures prior to passage. (E-F) Passage of culture using sterile syringes.

The remaining culture was stored in a closed CTB to simulate ambient storage on the ISS and return of live cultures to Earth (Figure 3A). At 36 d after the initial inoculation, all culture bags were sampled for DNA extraction and biomass assessment. The WT strain showed higher biomass compared to the *cw15* cell wall mutant, and there was a trend for increased biomass with additional culture passages. Increased biomass with the latter passages may reflect loss of biomass due to dark storage or selection for faster growth during the experiment.

**Figure 3.**
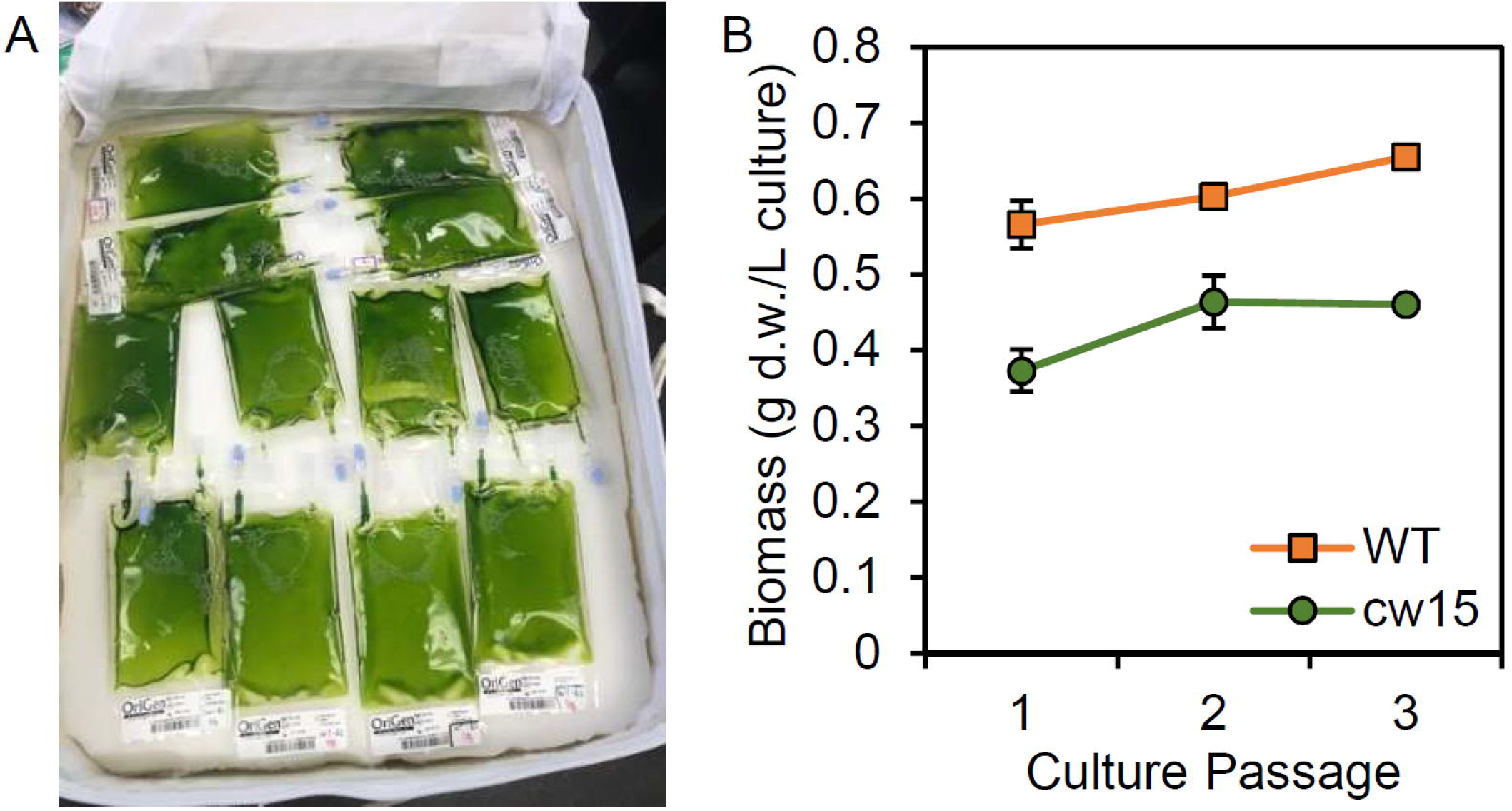
(A) Storage of live algae cultures in CTB soft stowage bag. (B) Biomass yield after dark storage. Average and standard deviation of three biological replicates are plotted.

Whole genome sequencing of the pre-mutagenized cultures, frozen cell pellets, and dark stored cultures was completed with an average read depth of 16x. Variants consisting of single nucleotide polymorphisms (SNP) and short insertion-deletion (InDel) polymorphisms were called with CRISP using pooled sample parameters (Figure 4). After removing missing data (≥ 30%) and monomorphic polymorphisms (MAF=0), we detected 73,573 WT variants and 79,455 *cw15* variants. Filtering to remove low-depth and low-quality reads reduced WT and *cw15* polymorphic variants to 48,380 and 53,939, respectively.

**Figure 4.**
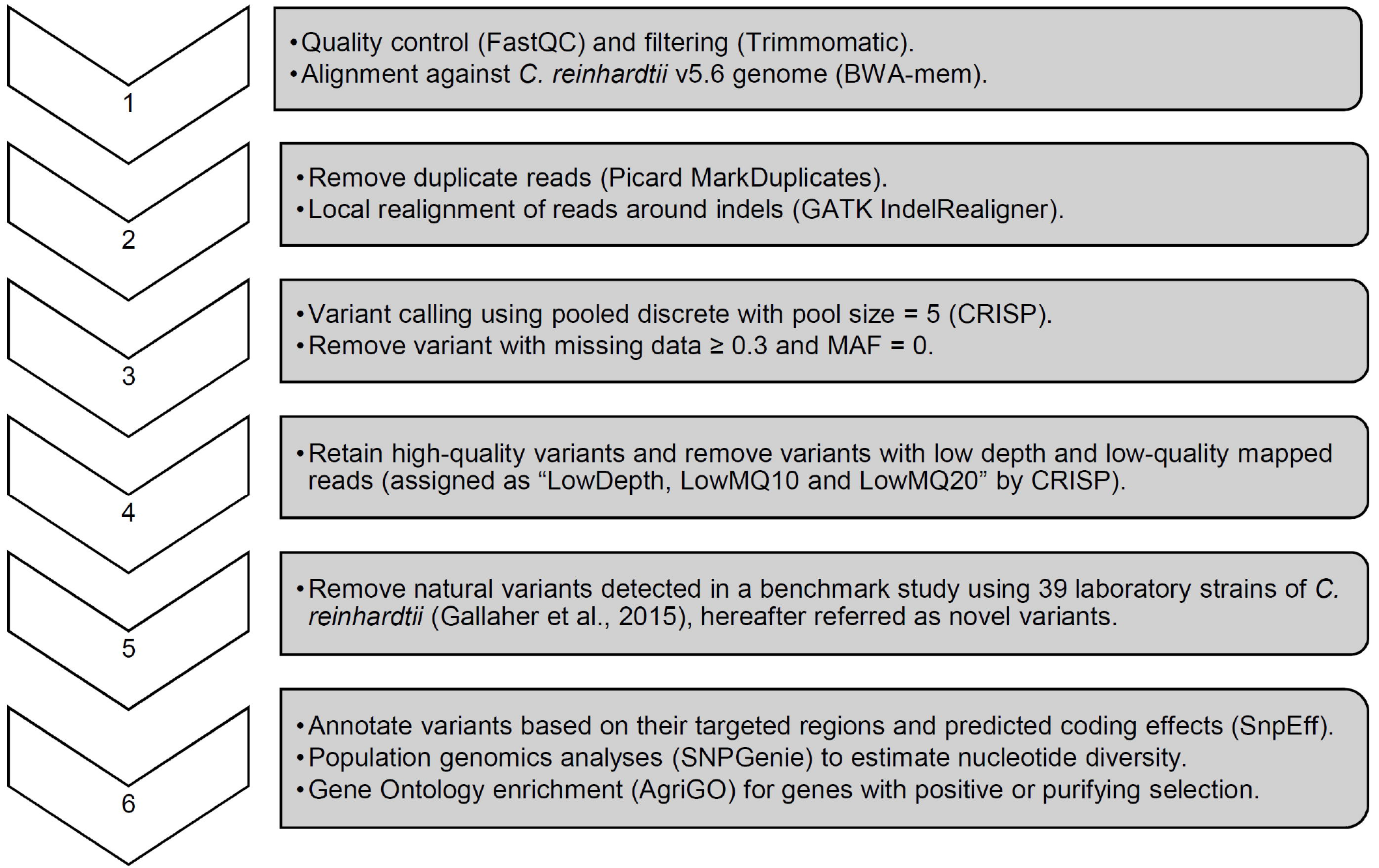
Sequence analysis pipeline with key parameters for variant calling and quality filtering.

Plotting the variant density across the genome identified known hotspots of natural variation among laboratory strains (Figure 5, Gallaher et al., 2015). The WT strain is a sequence-verified clone of CC-1690, and the polymorphic variants identified overlapping peaks on chromosomes 2, 6, and 9. In addition, there were peaks that overlapped with other natural variants on chromosomes 3 and 12. The *cw15* strain derives from a cross with CC-1690 and a similar pattern of natural variant peaks was observed. These natural variants may represent spontaneous mutations that are easily tolerated in Chlamydomonas or regions of the genome that are difficult to align with high confidence. In either case, known natural variants are likely to have little signal of selection due to induced UVC mutagenesis. We removed all exact matches to natural variants to obtain 5,286 WT and 5,873 *cw15* novel variants, which are more evenly distributed across the genome (Figure 5).

**Figure 5.**
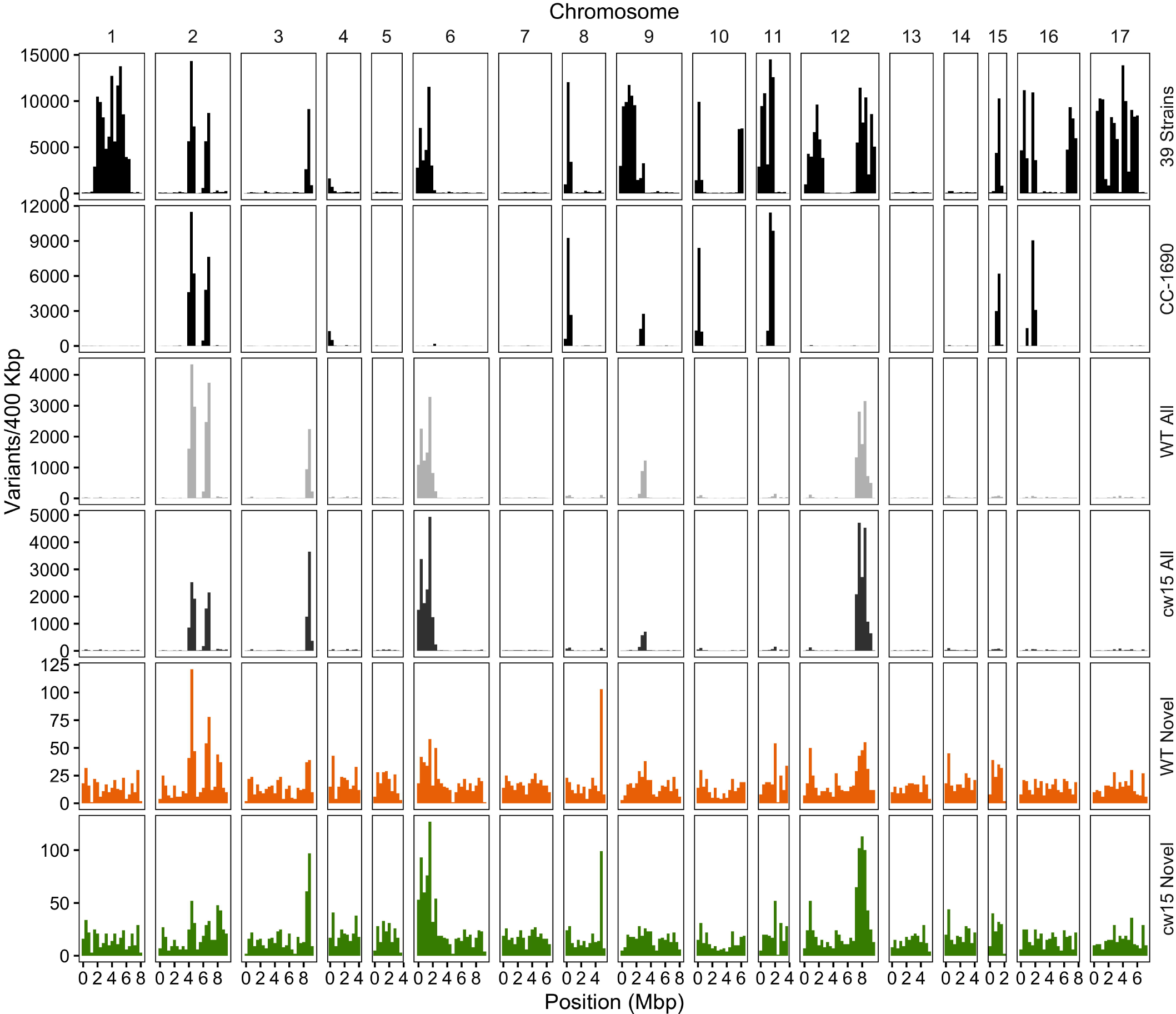
Density of SNP and indel variants in the Chlamydomonas genome. The Y-axis is the number of variants per 400 kb, and the X-axis is the physical distance of each chromosome. The variant data set plotted is labeled on the right of each panel. Variants from 39 laboratory strains (black, top panel) and CC-1690 (black, second panel) sequenced by Gallaher et al. (2015). Light and dark gray show all high quality variants (step 4 in Figure 4) in the WT (CC-5082) and *cw15* (CC-1883) strains. Orange (WT) and green (*cw15*) plot novel variants after filtering identical matches to polymorphisms in the Gallaher et al. (2015) study (step 5 in Figure 4).

The novel variants show a different spectrum of base changes than natural variants (Figure 6A). There is a relative decrease in transitions and an increase in transversions with the complementary mutations of A>C and T>G as well as C>G and G>C predominating. These data suggest the novel variants represent mutations caused by induced mutagenesis instead of the endogenous spectrum of Chlamydomonas variants. In addition, the novel mutations are found at a higher allele frequency indicating the novel mutations have more sequence read support across samples than natural variants (Figure 6B). We conclude that novel variants better represent the UVC mutagenized sites.

**Figure 6.**
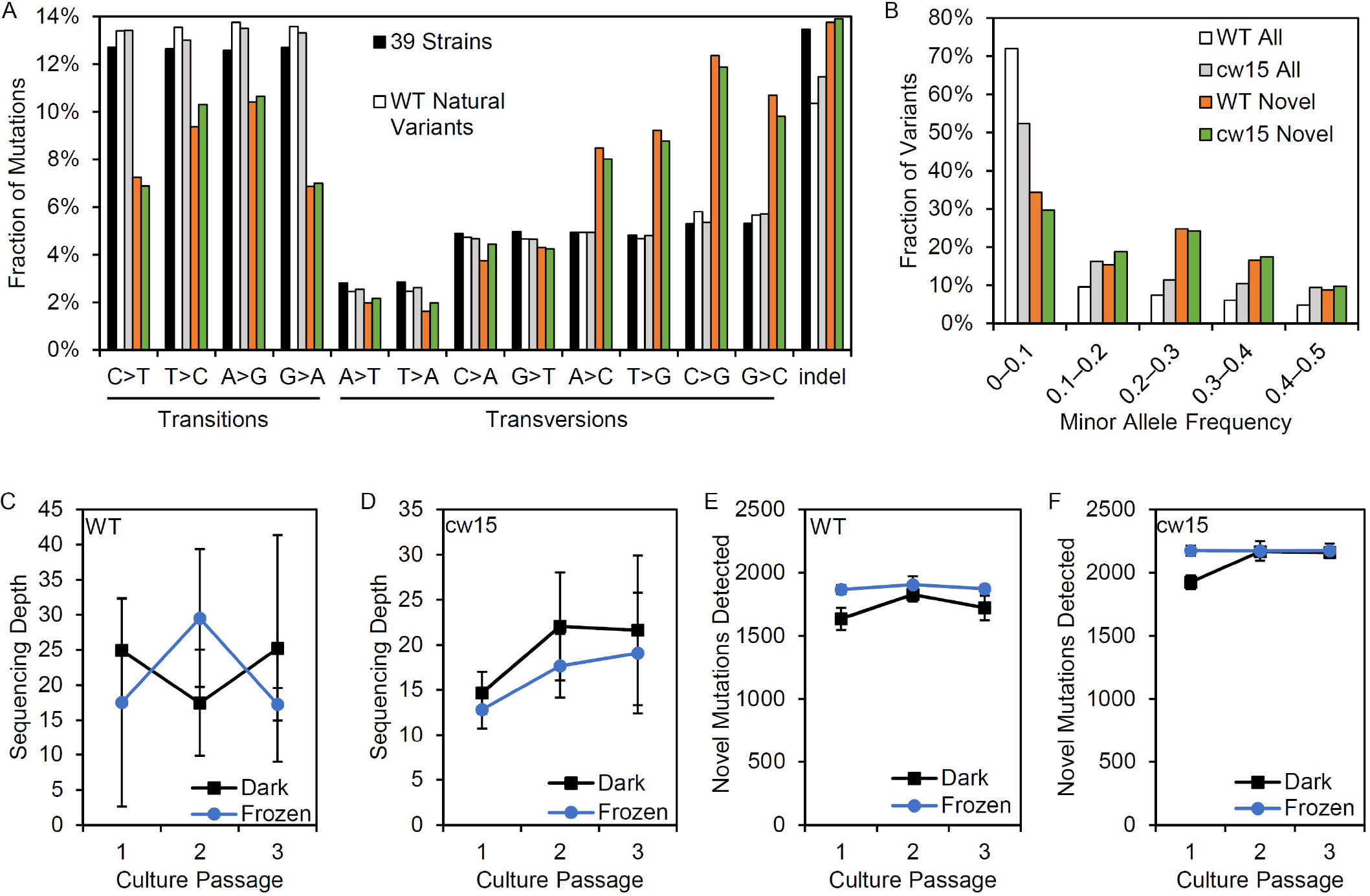
Effects of variant filtering and live culture storage on mutations detected. (A) Relative frequency of base changes and indels for natural variants found by Gallaher et al. (2015) and the novel variants after UV mutagenesis. (B) Minor allele frequency distribution for all variants and novel variants. (C-D) Average sequencing depth of three biological replicate libraries prepared from dark stored cultures and algae pellets that were frozen at the time of passage. (E-F) Average number of novel mutations detected in libraries from dark stored and frozen tissues. Error bars are standard deviations.

Centrifugation and freezing cell pellets for DNA sampling requires more extensive astronaut time and limiting resources on the ISS. We compared the mutations recovered from samples that had been frozen at the time of passage and those from live cultures that had been stored in the dark. There were no significant differences in sequencing depth based on the storage conditions (Figure 5C-D). However, dark storage decreased the number of mutations recovered in passage 1, which were cultures stored for 22 d prior to sampling. Student’s t-tests showed a significant reduction of mutations detected for the WT strain (p = 0.003), but the reduction was non-significant for *cw15* (p = 0.07). For frozen stored libraries, the number of novel mutations detected in each passage was nearly constant indicating sequencing depth was limiting for mutant detection. These results suggest that changes in allele frequency over culture passages is not a reliable indicator of selection for this experiment.

To assess the effects of novel mutation on protein coding sequences, SnpEff was used to identify protein coding changes. Natural variants were enriched for synonymous mutations, while the novel variants were enriched for protein coding changes (Figure 7A). These results are consistent with an increased frequency of deleterious mutations after UV mutagenesis. Individual genes were tested for selection based on nucleotide diversity (π) using SNPGenie. Novel variants were enriched for genes showing evidence of positive selection with about 46% of genes tested having π_N_ > π_S_, compared to 29% of all variants (Figure 7B). These results are consistent with the enrichment for nonsynonymous mutations resulting from UV mutagenesis.

**Figure 7.**
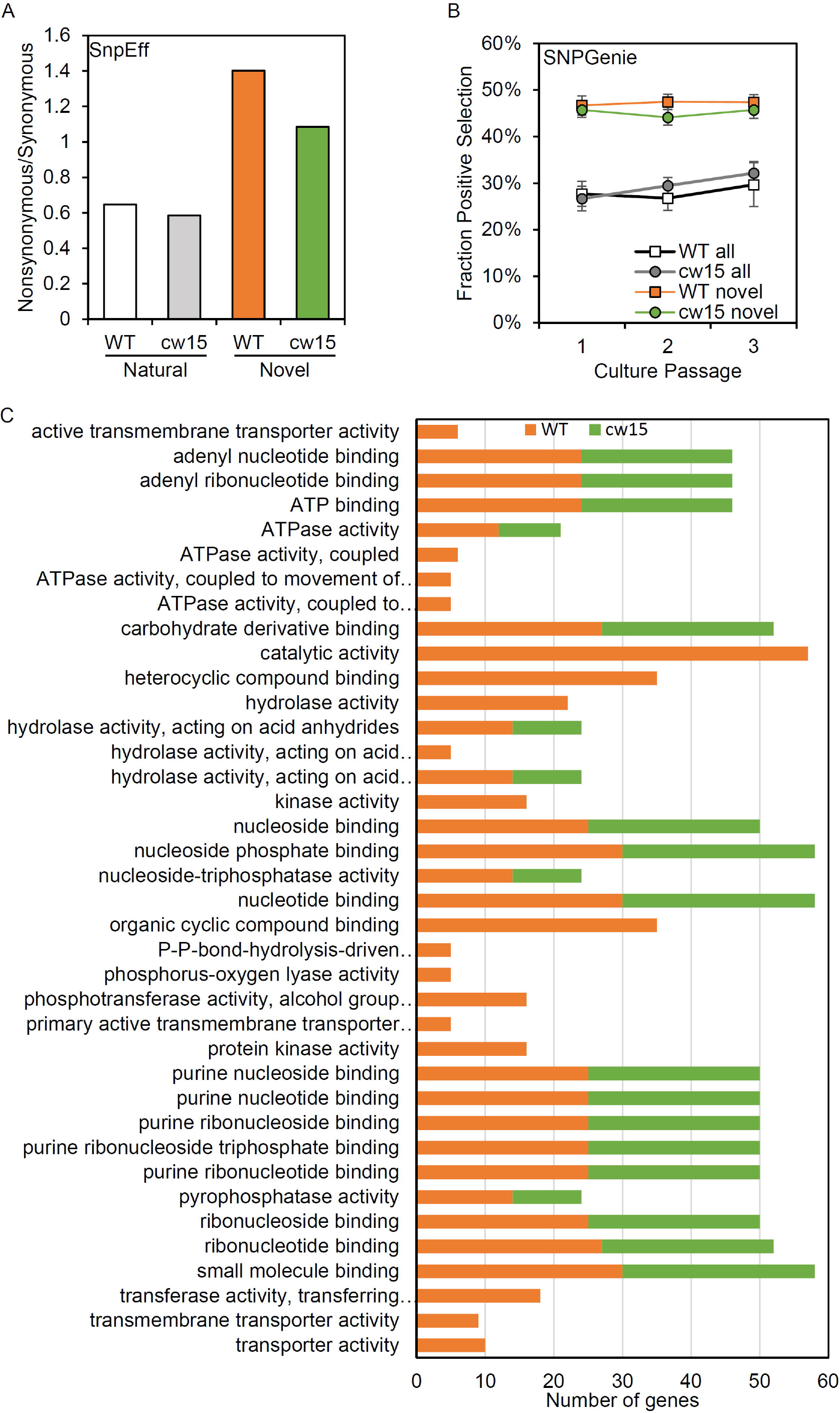
Novel variants are enriched for predicted protein coding changes. (A) Ratio of non-synonymous to synonymous variants based on SnpEff annotations. White and gray show natural variants detected. Orange and green show novel variants. (B) Average fraction of genes showing positive selection of all genes tested with SNPGenie. Averages are from six libraries per passage. Error bars are standard deviations. (C) Enriched GO terms for genes with positive selection.

Based on GO term enrichment analyses, the positively selected genes in WT and *cw15* both show significant enrichments for terms associated with purine nucleotide binding and hydrolase activity (Figure 7C). The individual genes with these GO terms represent information processing functions in DNA damage repair, RNA processing, translation, cytoskeletal motors, and signal transduction (Supplementary Table 1). The WT libraries also had enrichment for terms associated with small molecule transporters with individual genes predominantly being ABC transporters.

The WT strain also had significant enrichments for genes under purifying selection (Figure 8). More than half of these genes are predicted to function in regulating the levels of cyclic nucleotide second messengers suggesting second messenger signal transduction may be under selective pressure in in the culture bag growth system. The *cw15* strain had no significantly-enriched GO terms for genes under purifying selection.

**Figure 8.**
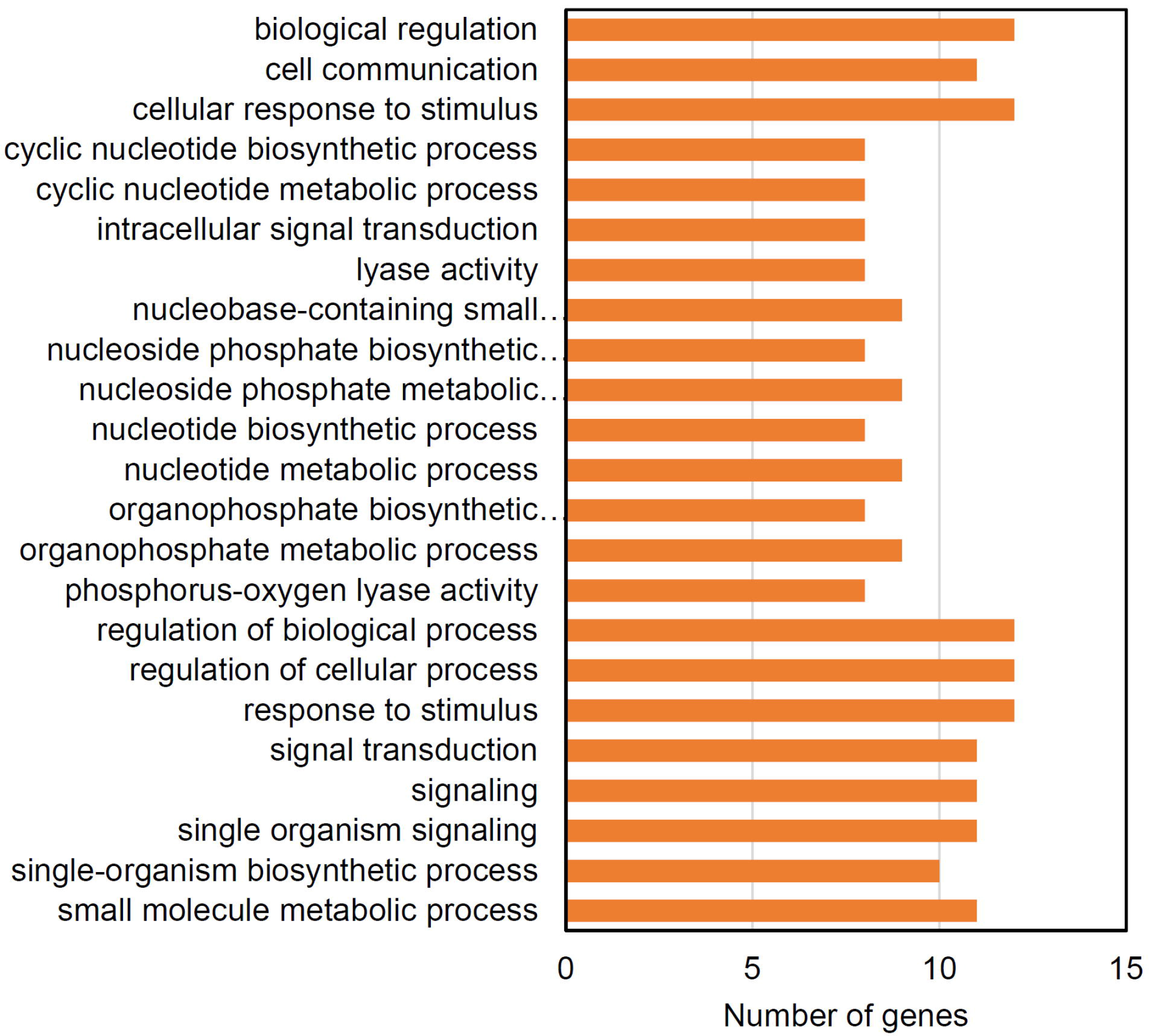
Enriched GO terms for genes showing evidence of purifying selection in the WT (CC-5082) strain.

## 4 Discussion

This EVT validated a strategy for identifying genes required by Chlamydomonas during log phase growth in spaceflight. We have shown that commercial FEP tissue culture bags can be used for batch culture of microalgae. Chlamydomonas is viable in these bags after prolonged dark storage, which enables full genome sequencing and identification of mutant genes in the culture. We have since used this strategy to grow the WT and *cw15* strains during the SpaceX CRS-15 mission; analysis of the spaceflight experiment is on-going.

Whole genome sequencing has been used to assess the mutagenic load of the bacteria, *Staphylococcus aureus*, in a two week spaceflight exposure (Guo et al., 2015). Less than 40 SNPs were detected in the genome from spaceflight with similar numbers of SNPs detected on ground and in spaceflight. These data suggest that exogenous mutagenesis is necessary to gain adequate signal of selection in short-term competitive growth experiments.

A yeast selection experiment was completed in spaceflight by using a genome-wide deletion collection (Nislow et al., 2015). The competitive growth experiment measured the reduction in representation of bar-coded mutants over the course of ~21 mitotic generations. This type of competitive growth identifies individual genes needed for growth. By contrast, UVC mutagenesis has a higher genetic load and generates a more diverse array of allele types to compete within the culture. Limitations to random mutagenesis are that loss-of-function mutations in all genes are not represented in each biological replicate of the experiment and that multiple mutations are simultaneously selected in individual cells during mitotic divisions.

UV light causes direct DNA damage to create cyclobutane pyrimidine dimers, and the UV mutation signature is typically biased towards C>T transitions (Ikehata and Ono, 2011). Chlamydomonas DNA readily forms pyrimidine dimers, and WT strains have a robust dark-repair pathway to repair 90-95% of the DNA damage directly within 24 h (Small, 1987). With whole genome sequencing, we observed enrichment for T>G and C>G transversions in the mutations resulting from the UV treatment instead of the expected C>T transitions. Incorporation of several oxidative products of guanine can promote C>G transversions in response to a variety of mutagens, including UV (Kino and Sugiyama, 2005). In human cancers, C>G base changes are associated with activation of AID/APOBEC cytidine deaminases followed by error-prone translesion synthesis (Forbes et al., 2017). Human cancers also have base substitution signatures that are enriched for T>G, such as COSMIC signatures 17b and 28 (Forbes et al., 2017). However, no specific mechanisms have been proposed for the specific enrichment of T>G transversions.

Intriguingly, we observed signatures of selection in DNA polymerase ζ, θ, and REV1 (Supplementary Table 1), which are associated with translesion synthesis (Sakamoto, 2019). In the human germ line, C>G mutations have been suggested to be caused by errors in double-stand break repair (Gao et al., 2019). DNA polymerase θ and DNA ligase 4 function in the non-homologous end joining double strand break repair pathway, and we found evidence of selection for both of these enzymes in the EVT (Pannunzio et al., 2018). Evidence for selection in double-strand break repair and translesion synthesis suggest that these pathways are likely relevant to the UV-induced mutations observed.

The low sequence coverage of this experiment creates risk in using the frequency of recovery for specific mutant alleles in determining whether specific genes are under positive or purifying selection. The number of alleles discovered is limited by the sequencing depth, and higher coverage is necessary to increase the power of the statistics to detect selection at a genome-wide level. Nevertheless, we were able to classify 476 genes in the WT strain and 503 genes in the *cw15* strain for purifying or positive selection based on mutations within coding sequences. Among these, 104 genes were enriched for molecular functions based on GO terms. In addition to DNA repair pathways, these analyses revealed signal transduction and other information processing functions including, chromatin reading, RNA processing, and translation to be enriched within selected genes. The enriched molecular functions are predicted to be processes necessary for Chlamydomonas to adapt to the UV mutagenesis, lack of media agitation, the diffusion of gases across the FEP membrane, and the lighting conditions in the Veggie unit.

Scalable production of Chlamydomonas in spaceflight has multiple potential applications. The species will accumulate lipids to about 20-25% of total biomass, which can be used as an organic chemical feedstock (Becker, 2007; Xu et al., 2018). Chlamydomonas also has high protein content of 40-60% of biomass making it a potential source of food. Although it is not yet designated as a GRAS organism by the FDA, Chlamydomonas is non-toxic; animal feeding studies show no harmful effects at 4 g algae biomass per kg body weight per day, the highest consumption levels tested (Murbach et al., 2018). Equivalent consumption for a 65 kg person would be approximately 260 g of algae powder per day.

As a genetic model organism, Chlamydomonas biomass composition can be further modified by mutagenesis or targeted gene editing. For example, starch over-accumulation mutants have been isolated that shift starch content from 15% to 30-35% of biomass (Koo et al., 2017). Likewise, CRISPR gene editing of the Chlamydomonas zeaxanthin epoxidase gene significantly increases carotenoids needed to prevent macular degeneration (Baek et al., 2018). The modified algae strain was used to supplement chicken feed to increase the zeaxanthin content of eggs. The myriad of potential applications for microalgae production in spaceflight justify direct investigation of the genes needed for liquid culture production. Our current study shows both feasibility for a spaceflight experiment and identifies a series of cellular information processing genes that are likely required for Chlamydomonas to adapt to batch culture in breathable plastic bags.

## 5 Data Availability Statement

The whole-genome sequences generated for this study can be found in the GeneLab database. https://genelab-data.ndc.nasa.gov/genelab/accession/GLDS-265/

## 6 Acknowledgments

We thank Matthew Hoffman and Nicole Dufour for organizing the EVT at the Kennedy Space Center Veggie facility. This work was funded by the Center for the Advancement of Science in Space (currently International Space Station U.S. National Laboratory), the Florida Space Institute and the Vasil-Monsanto Endowment.

## 7 Author Contributions Statement

JZ, BR, KT, FB, and AS developed and performed the batch culture and mutagenesis. JZ constructed whole genome sequencing libraries and preliminary processing of the data. BM, YH, MR, and AS completed genome sequence analysis and interpretation. AS, JZ, and BM wrote the manuscript. All authors reviewed and agree with the manuscript content.

## 9 Supplementary Material

The Supplementary Material for this article can be found online at:

## 10 Contribution to the Field Statement

Microalgae are single-celled, photosynthetic organisms that can be grown to capture carbon dioxide and recycle nitrogenous wastes. Most algae species require low levels of light and are highly efficient at carbon capture. Since nearly the inception of international space programs, microalgae have been proposed to be part of biological regenerative support systems for long-duration spaceflight. Scalable production systems for growing algae in liquid cultures have not been developed for microgravity conditions. Moreover, the genetic adaptations needed to maximize carbon capture during spaceflight by microalgae species have not been determined. We developed a simple protocol for batch liquid cultures using commercial breathable plastic bags. This system can easily scale in volume depending upon the size of available growth facilities. We also established and tested a genetic selection experiment to identify genes required for the model algae, *Chlamydomonas reinhardtii*, to grow in the plastic bag cultures in the Veggie plant growth chamber. These ground-based verification experiments show that selected for improved microalgae productivity in space is feasible.

